# Learning-dependent feedback by OLM interneurons shapes CA1 representations

**DOI:** 10.64898/2025.12.21.695825

**Authors:** Evan P. Campbell, Lisandro Martin, Jeffrey C. Magee, Christine Grienberger

## Abstract

Spatial learning depends on hippocampal CA1 place cell representations, which form rapidly through behavioral timescale synaptic plasticity (BTSP). BTSP is driven by dendritic plateau potentials proposed to arise from the interaction of an excitatory target signal from entorhinal cortex layer 3 (EC3) and inhibitory feedback reflecting the current CA1 population state. However, the cellular source of this feedback has remained unknown. Using two-photon Ca^2+^ imaging in mice during spatial learning, we found that dendrite-targeting oriens–lacunosum moleculare (OLM) interneurons increased their activity at behaviorally salient locations in a manner consistent with previously described environment-specific CA1 representations and EC3 target signals. Causal manipulations revealed that silencing a genetically defined subset of OLM interneurons late in learning enhanced BTSP and place field formation, whereas activating them early suppressed place field formation. These findings identify OLM interneurons as a key inhibitory feedback element regulating BTSP and the formation of hippocampal representations during learning.

## Introduction

Learning is accompanied by persistent, experience-driven changes in neural activity that support the acquisition and recall of behaviors critical for survival. Newly learned information is thought to be stored in coordinated activity patterns and synaptic weights within neuronal ensembles, which are reactivated during memory retrieval^1-5^. Spatial learning engages behavioral timescale synaptic plasticity (BTSP) in hippocampal area CA1, which is triggered by Ca^2+^ plateau potentials (“plateaus”) in the distal apical tuft dendrites of CA1 pyramidal neurons^6-8^ and generates CA1 place cell representations^9-15^. In previous work, we showed that entorhinal cortex layer 3 (EC3) provides a stable, target-like excitatory input that is enhanced at behaviorally salient locations and promotes dendritic plateau initiation^12^, thereby driving BTSP and shaping CA1 population activity toward an overrepresentation of salient locations required for successful spatial learning^16-20^. Notably, this EC3 target signal remains stable across learning, whereas BTSP-dependent place field formation is strongest early and declines with experience^12^. We therefore propose a model in which plateau initiation reflects a mismatch between the EC3-derived excitatory target signal and an inhibitory feedback signal reporting the current state of CA1 population activity. In this framework, plateaus emerge when excitatory drive exceeds feedback inhibition (Fig. 1a). However, the cellular origin of this feedback remains unknown. Oriens–lacunosum moleculare (OLM) interneurons are well positioned to provide such feedback, as they receive excitatory input from CA1 pyramidal neurons and selectively target the tuft dendritic compartment^21-23^, where plateau initiation occurs.

**Fig. 1.**
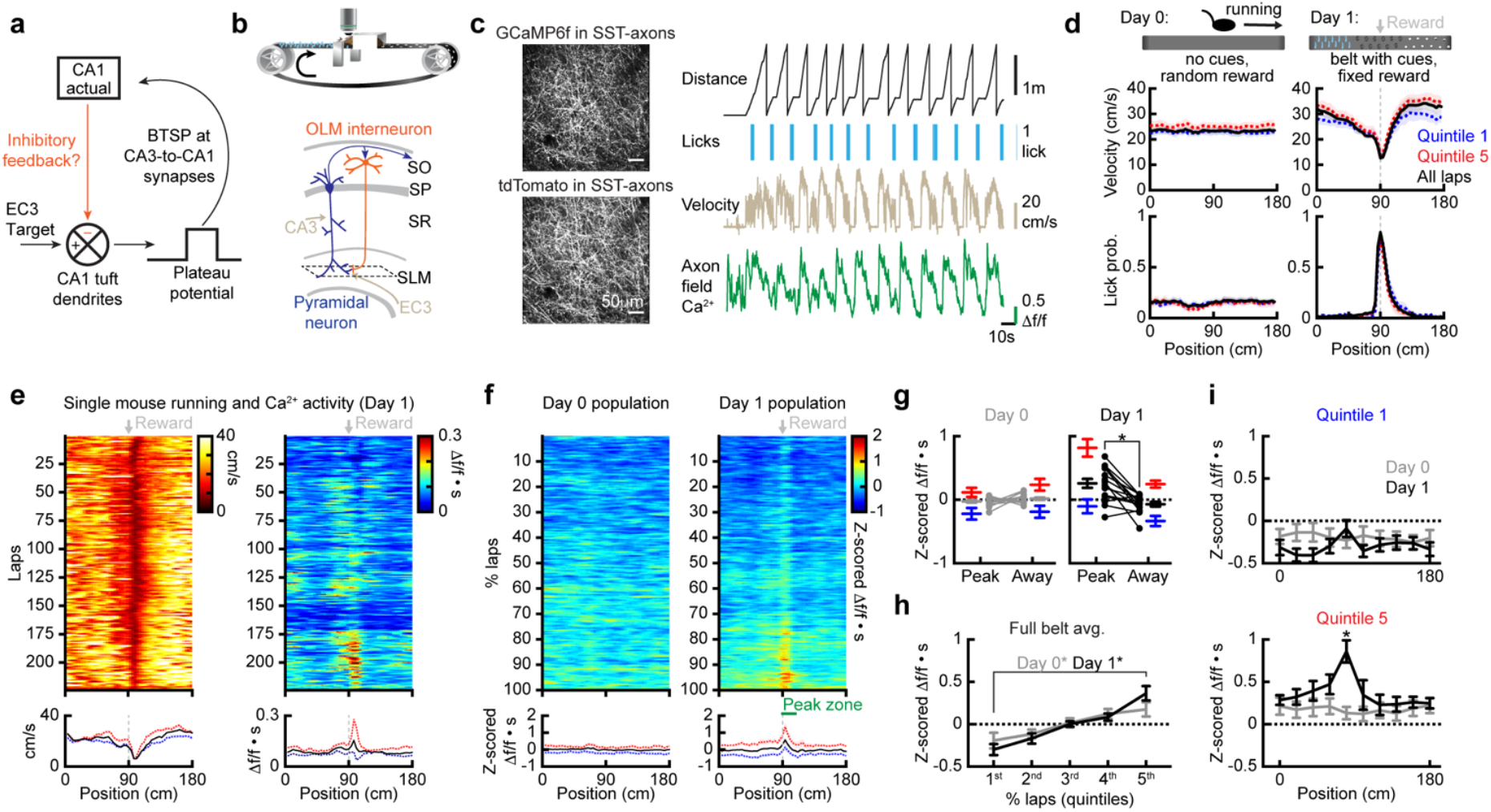
OLM population activity during learning. **a**, BTSP working model. **b**, Top: experimental setup, in which mice learn the location of a water reward. Bottom: schematic of the CA1 microcircuit and imaging plane (dashed line) **c**, Left: representative, two-photon, time-averaged images of axons from CA1 SST-OLMs expressing GCaMP6f and tdTomato. Right: example traces of distance, licking, velocity, and GCaMP6f Δf/f (mean of the field). **d**, Top: Task design of day 0 (the last day of habituation, no cues, random reward), and day 1 (the first day of learning, belt with tactile cues and fixed reward. Middle: mean running profile across space. Bottom: mean lick probability across space (blue: quintile 1, red: quintile 5, black: all laps). **e**, Representative day 1 running, and Δf/f corrected for dwell time (•s). Top: along laps. Bottom: spatial profiles. **f**, Days 0 and 1 mean z-scored Δf/f•s. Top: along the percentage of laps run. Bottom: spatial profiles. The peak-zone is 18 cm beginning at the spatial bin with reward, the away-zone is the same but 90 cm away. **g**, Mean z-scored Δf/f•s within peak-zone and away-zone (blue: quintile 1, red: quintile 5, black/gray: all laps). Left: day 0. Right: day 1 (paired two-tailed *t*-test, P=4•10^-4^). **h**, Mean z-scored Δf/f•s vs. lap quintiles (two-way RM ANOVA with post-hoc FDR correction, day 0/quintile 1 vs. 5: P=5•10^-4^, day 1/quintile 1 vs. 5: P<1•10^-4^). **i**, Quintiles 1 (top) and 5 (bottom) mean z-scored Δf/f•s vs. position (bin=18 cm, two-way RM ANOVA with post-hoc FDR correction, quintile 1/group x bin: P=<1•10^-4^, quintile 5/group x bin: P<1•10^-4^, quintile 5/bin 5: P<1•10^-4^). In **d**,**f**-**i**, n=15 mice. In **g**, dots represent mice. Data are shown as mean ± s.e.m.

Here, we combined in vivo two-photon Ca^2+^ imaging with bidirectional optogenetic manipulations to examine the role of OLM interneurons in spatial learning and their function within the CA1 microcircuit shaping BTSP-dependent representations. Axonal and somatic OLM activity increased with learning and was biased toward behaviorally salient locations, mirroring the emergence of CA1 population representations aligned with the environment-specific EC3 target signal. Optogenetic silencing of Chrna2α-positive OLM interneurons late in learning revealed enhanced BTSP induction and place field formation in pyramidal neurons, specifically at locations where the EC3 signal is high^12^. Conversely, optogenetic activation of Chrna2α-positive OLM interneurons early in learning suppressed place field formation at stimulated locations. Together, these findings demonstrate that OLM interneurons provide learning-dependent inhibitory feedback that regulates BTSP induction and the formation of CA1 spatial representations.

## Results

### OLM axonal activity increases with spatial learning

To determine whether OLM interneurons constitute the inhibitory feedback signal involved in the BTSP model, we first performed two-photon Ca^2+^ imaging of OLM axons in dorsal CA1 of somatostatin (SST)-IRES-Cre mice^24^ while animals learned a fixed reward location (Fig. 1b). OLM axonal Ca^2+^ signals were recorded in *stratum lacunosum-moleculare* (SLM), the CA1 sublayer where entorhinal cortex layer 3 (EC3) inputs converge with OLM-mediated inhibition, and Ca^2+^ signals from all axons within a field of view were averaged to yield a population-level measure of OLM output alongside behavioral variables (Fig. 1c, Extended Data Fig. 1a). As in prior work^12,16^, when learning the location of a fixed reward (day 1, belt with tactile cues, fixed reward), animals confined licking to the reward vicinity and slowed during reward approach, as revealed by comparing early (quintile 1, blue) and late laps (quintile 5, red; Fig. 1d). These behavioral changes suggest a learning of the reward location and were absent during the day 0 sessions (last day of habituation; no cues, random reward). After correcting for dwell time to account for learning-related changes in running speed^25-27^ (see Method section), OLM axonal Ca^2+^ activity on day 1 and subsequent sessions developed spatial structure with learning, showing a modest global increase across the track and a pronounced peak at the reward location during later laps, whereas the day 0 dwell time-corrected OLM activity increased uniformly without spatially localized modulation (Fig. 1e–i, Extended Data Fig. 1b–g). A similar learning-dependent increase in OLM activity was observed in a distinct environment in which animals ran on a blank belt, with a visual cue positioned 50 cm before the reward. Under this condition, activity also increased globally but peaked near the reward-predictive cue location (Extended Data Fig. 2).

In our prior work, we showed that EC3 target patterns differ across these two environments and shape the resulting CA1 population representations^12^. Here, we find that OLM activity differs across these environments in a manner consistent with these previously described changes in CA1 representations. These results suggest that OLM interneurons track the state of CA1 population activity during learning and, thus, identify OLM interneurons as a potential candidate source of the learning-dependent inhibitory feedback signal that interacts with the entorhinal excitatory input at distal apical dendrites to regulate plateau initiation. Notably, we observed the OLM activity development despite averaging across all axons within the field of view, potentially suggesting a population-level modulation of OLM output rather than the emergence of highly selective individual axons.

### Spatial learning induces population-wide changes in OLM interneuron activity

To determine whether the learning-related OLM activity changes observed at the population axonal level reflected changes in many OLM interneurons rather than being driven by a small subset of strongly modulated cells, we examined Ca^2+^ dynamics in the somata of identified OLM interneurons during the tactile cue paradigm. For these experiments, we used the Chrna2α-Cre mouse line, which provides selective and reliable genetic access to OLM interneurons^22,28^, and performed two-photon Ca^2+^ imaging of OLM somata located in *stratum oriens* (Fig. 2a). Example traces from three OLM neurons illustrate heterogeneous but coordinated activity across laps (Fig. 2b, h). We first quantified the population-averaged OLM somatic activity (Fig. 2c-g, see Methods). We found that it closely matched the spatial and temporal profiles of SST axonal activity (Fig. 1), indicating that the axonal signals reflect changes in OLM activity rather than additional axonal processing. Next, we examined how learning shaped activity patterns at the individual neuron level (Fig. 2h-o). Spatially binned, dwell time-corrected activity maps from three example neurons illustrate an increase in activity primarily at the reward location over the course of the day 1 session in all three cells (Fig. 2h). Consistently, heatmaps summarizing the activity of all recorded Chrna2α-positive OLM neurons (n=57) during early (quintile 1) and late (quintile 5) learning reveal a population-wide increase in activity at the reward location (Fig. 2i). In line with this observation, quantitative analyses showed modest but significant increases in spatial and reward selectivity in quintile 5 compared to quintile 1 across the OLM population (Fig. 2j–l). In addition, the peak activity location of individual neurons shifted toward the reward location over the course of the session (on average 13.3 +/-3.1 cm), with a wide distribution of shift magnitudes, but with 73.7% of the population shifting closer to reward (Fig. 2m). Notably, neurons whose activity peaks were initially located farther from the reward exhibited larger shifts by the end of learning (Fig. 2n). Finally, pair-wise correlation between neurons recorded in the same animal also increased (Fig. 2o). Together, these analyses establish that learning-related changes in OLM activity are distributed across the population, with most neurons contributing by varying degrees to the evolving increase in inhibitory feedback signal.

**Fig. 2.**
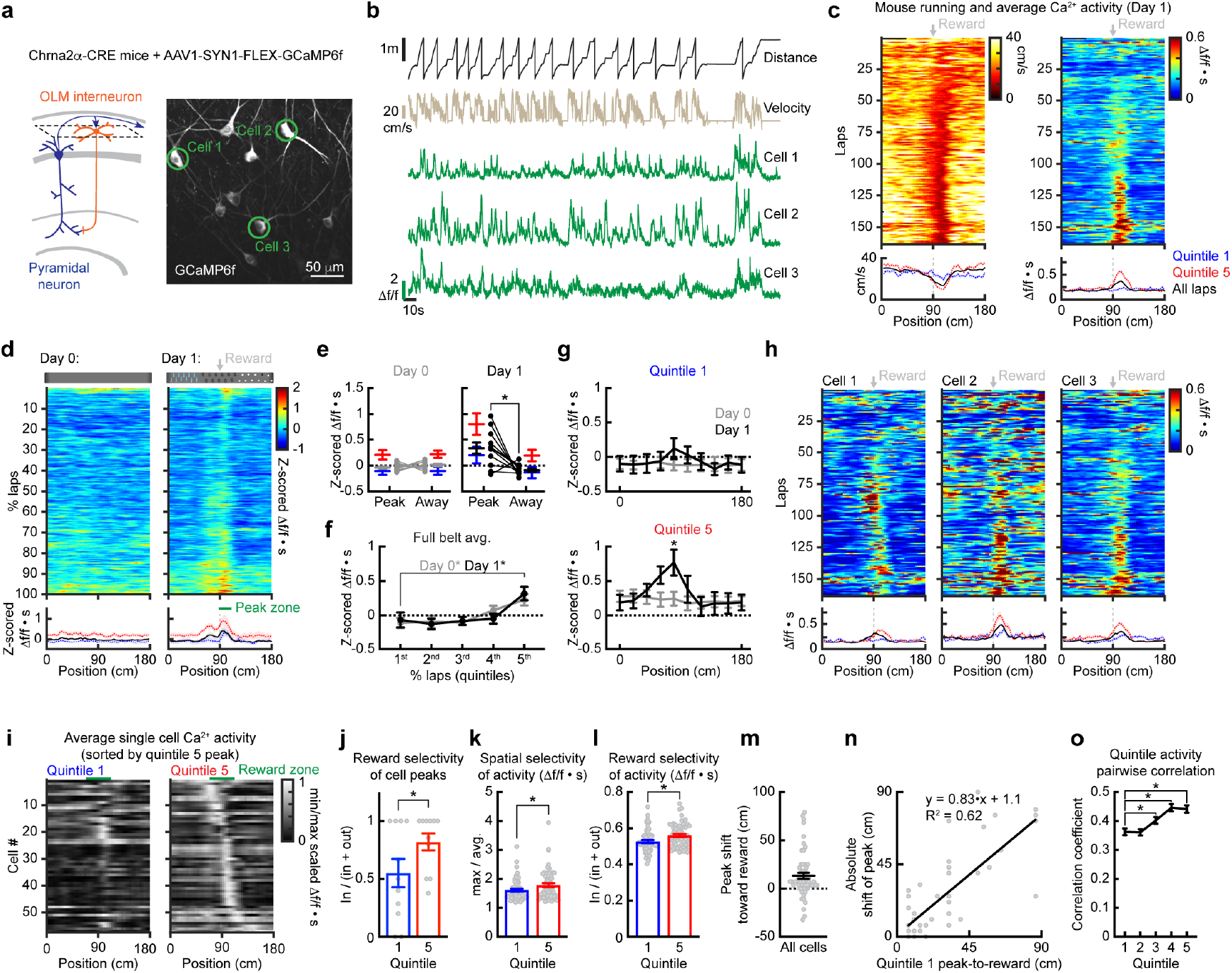
Single-cell OLM activity during learning. **a**, Left: Schematic of the CA1 microcircuit with imaging plane (dashed line). Right: representative, two-photon, time-averaged image of Chrna2Δ-OLMs expressing GCaMP6f, corresponding to **b**,**c**,**h. b**, Example traces of distance, velocity, and Δf/f for three OLM neurons. **c**, Representative day 1 running and mean Δf/f•s. Top: along laps. Bottom: spatial profiles (blue: quintile 1, red: quintile 5, black: all laps). **d**, Top: Task design of day 0 and day 1. Middle: mean z-scored Δf/f•s along the percentage of laps run. Bottom: spatial profiles. The peak-zone is 18 cm beginning at the spatial bin with reward, the away-zone is the same but 90 cm away. **e**, Mean z-scored Δf/f•s within the peak-zone and away**-**zone (blue: quintile 1, red: quintile 5, black/gray: all laps). Left: day 0. Right: day 1 (paired two-tailed *t*-test, P=8.8•10^-3^). **f**, Mean z-scored Δf/f•s vs. lap quintiles (Two-way RM ANOVA with post-hoc FDR correction, day 0/quintile 1 vs. 5: P=3.4•10^-2^, day 1/quintile 1 vs. 5: P=1.1•10^-2^). **g**, Quintiles 1 (top) and 5 (bottom) mean z-scored Δf/f•s vs. position (Two-way RM ANOVA with post-hoc FDR correction, quintile 5/group x bin: P=7•10^-4^, quintile 5/bin 5: P=1.8•10^-3^). **h**-**o**, Day 1 recordings of individual Chrna2Δ-OLMs. **h**, Δf/f•s for three representative neurons. Top: along laps. Bottom: spatial profiles. **i**, Min/max scaled mean Δf/f•s for all neurons (n=55 neurons, scaling from quintiles 1 and 5, sorted by quintile 5 peak). The reward zone is 39.6 cm centered at the reward. **j**, Mouse reward selectivity (the fraction of neurons with mean Δf/f•s peaks within the reward-zone) (paired two-tailed *t*-test, P=2.0•10^-2^). “In” and “out” refer to cells that have peaks inside or outside the reward zone. **k**, OLM spatial selectivity (max/mean Δf/f•s) (paired two-tailed *t*-test, P=4.4•10^-2^). **l**, OLM reward selectivity (the fraction of mean Δf/f•s within the reward-zone) (paired two-tailed *t*-test, P=1.1•10^-2^). “In” and “out” refer to activity inside or outside the reward zone. **m**, Mean distance that Δf/f•s peaks shifted toward reward from quintile 1 to 5. **n**, The absolute distance Δf/f•s peaks shifted from quintile 1 to 5, vs. the absolute peak-to-reward distance in quintile 1. **o**, Pooled pairwise Pearson’s correlation coefficients from co-recorded neurons vs. quintile (n=166 unique pairs, one-way RM ANOVA with post-hoc FDR correction, P<1•10^-4^, quintiles 1/3 P=2•10^-4^, quintiles 1/4 P<1•10^-4^, quintiles 1/5 P<1•10^-4^). In **d**-**f**,**i**-**o** Day 0: n=12 mice, Day 1: n=11 mice. In **e**,**j** dots represent mice, and in **k**-**n** they represent neurons. Data are shown as mean ± s.e.m.

### Silencing OLM interneurons late in learning permits BTSP-mediated place cell formation

If OLM activity indeed constitutes the inhibitory feedback predicted to regulate BTSP, then manipulating OLM activity should bidirectionally constrain plateau initiation and place field formation. To test this hypothesis, we first used a combined transgenic and viral strategy to perform Ca^2+^ imaging of GCaMP6f-expressing CA1 pyramidal neurons while optogenetically suppressing OLM interneuron activity via archaerhodopsin-T (ArchT)^29^ during spatial learning (Fig. 3a-b). The optogenetic manipulation was restricted to an “opto-zone” centered around the fixed reward location (Fig. 3a). Each session was divided into two blocks, with no light delivered during block 1 (laps 1–50) and OLM silencing via 594 nm light during block 2 (laps 51–100). This design allowed us to perturb OLM activity late in learning around the reward, when and where OLM activity is highest (Figs. 1–2). Mice that did not express ArchT but were otherwise treated identically served as controls. In a subset of animals, we confirmed that 594 nm-light effectively suppressed the activity of ArchT-expressing Chrna2α-positive interneurons (Extended Data Fig. 3a–f). Importantly, this optogenetic manipulation did not alter running or licking behavior, indicating that ArchT activation did not disrupt task engagement or reward anticipation (Fig. 3c). Consistent with the absence of optogenetic manipulation during block 1, CA1 place cell activity was similar in control and ArchT-expressing animals during this phase of the session (Fig. 3d–e, Extended Data Fig. 4a–c). In contrast, silencing OLM neurons during block 2 resulted in an increase in place cell density within the opto-zone around the reward in ArchT-expressing animals compared with controls (Fig. 3d–e). To quantify the temporal dynamics of this effect, we analyzed the emergence of new place cells, distinguishing cells whose place field peaks fell within the opto-zone from those located in an “away-zone” approximately 90 cm from the opto-zone (Fig. 3f). In ArchT-expressing animals, new place field formation increased within the opto-zone during block 2, with no corresponding change in the away-zone. Consequently, the fraction of CA1 place cells was significantly higher in the opto-zone during block 2 in ArchT-expressing animals compared to controls, while no differences were observed in block 1 or in a same-sized away-zone (Fig. 3g). Together, these data indicate that OLM suppression selectively facilitates place cell formation at the manipulated location and late stage of learning.

**Fig. 3.**
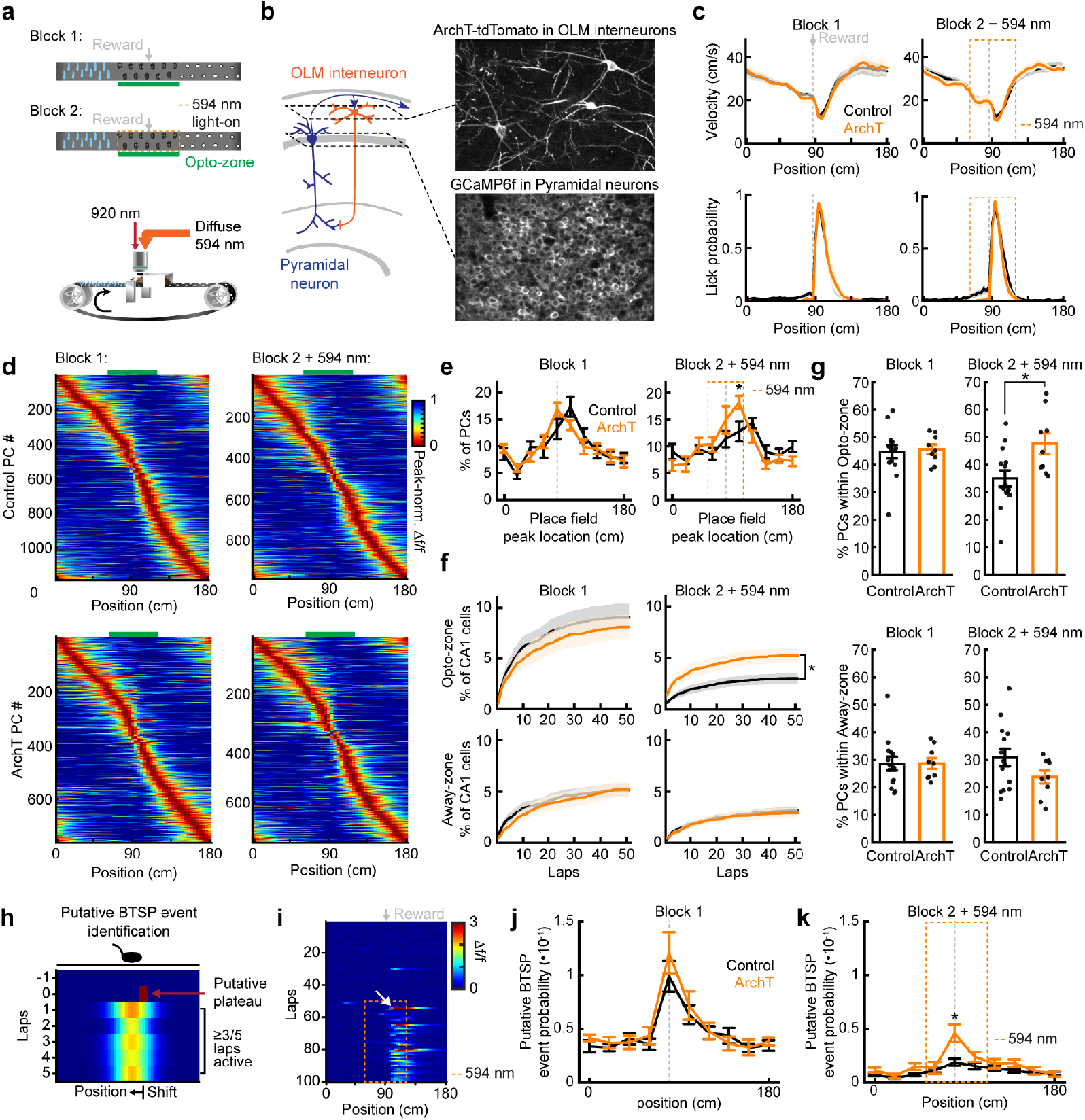
Silencing Chrna2Δ-OLMs late in learning. **a**, Top: day 1 manipulation paradigm wherein ArchT in Chrna2Δ-OLMs is activated during block 2 (50-lap blocks) within 30 cm of the reward (the opto-zone is 60 cm, centered at the reward, the away-zone is the same, but 90 cm away from the opto-zone, orange dashed box depicts 594 nm-light on period). Bottom: Imaging and optogenetics apparatus. **b**, Left: Schematic of the CA1 microcircuit with two imaging planes (dashed lines). Right: representative, two-photon, time-averaged images of Chrna2Δ-OLMs expressing Arch-tdTomato and pyramidal neurons expressing GCaMP6f. **c**, Blocks 1 and 2 behavior. Top: mean running profile across space. Bottom: mean lick probability across space. **d**, Blocks 1 and 2 peak-normalized mean Δf/f vs. position for all place cells (PCs), analyzed independently within blocks, sorted by place field peak location. Top: control (block 1: n=1188, block 2: n=989). Bottom: ArchT (block 1: n=762, block 2: n=748). **e**, Blocks 1 and 2 fractions of CA1 place cells vs. place field peak location (unpaired two-tailed *t*-test: block 2/bin 5 P=4•10^-2^). **f**, Blocks 1 and 2 CA1 place cell onsets. Top: place cells with place field peaks within the opto-zone (two-tailed *t*-test, block 2: P=1•10^-2^). Bottom: same for away-zone. **g**, Place cell distributions in the opto-zone and away-zones. Left: block 1, Right: block 2 (unpaired two-tailed *t*-test, opto-zone: P=1•10^-2^). **h**, Schematic of criteria for putative BTSP events. **i**, Example place cell from an ArchT mouse. Mean Δf/f along laps with putative BTSP event marked by the white arrow. **j**. Block 1 mean putative BTSP event probability for all cells vs. position. **k**, Same as **j**, but block 2 (two-way RM ANOVA with post-hoc FDR correction, group x bin: P=7•10^-4^, bin 5: P=0.012). In **c**-**g**,**j**-**k**, control: n=14 mice, ArchT=9 mice. In **g** dots indicate mice. Data are shown as mean ± s.e.m.

To determine whether silencing OLM interneurons in the block 2 opto-zone was also associated with an increased incidence of BTSP events, we next identified putative BTSP events using established GCaMP6f imaging signatures derived from prior whole-cell and imaging studies^5,9,12,30^. Putative BTSP events were defined as large-amplitude Ca^2+^ transients followed by activity across at least three of five subsequent laps that shifted forward relative to the event, with only the first qualifying event per cell included in the analysis (Fig. 3h–k, Extended Data Fig. 5a-b, see Method section). We found that the CA1 population in control animals exhibited a marked reduction in putative BTSP event probability from block 1 to block 2 (Fig. 3j–k; 46% ± 4 of active CA1 neurons exhibited an event in block 1 vs. 10% ± 0.8 in block 2, full belt, mean ± SEM, paired two-tailed *t-*test, P<1•10^-4^), with events preferentially concentrated near the reward location. An overall reduction was also observed in ArchT-expressing animals (51% ± 4 vs. 17% ± 1, block 1 vs. block 2 full belt; paired two-tailed t-test, P<1•10^-4^). However, during block 2, the probability of observing a putative BTSP event was significantly higher in ArchT-expressing animals than in controls (unpaired two-tailed t-test, *P*<1•10^-4^). This effect was driven by an approximately 2.5-fold increase in the probability of putative BTSP events within the center of the opto-zone relative to controls (Fig. 3j–k), indicating a selective enhancement of BTSP activity at the manipulated location. Notably, the amplitudes of the lower 80% of Ca^2+^ events (used as a proxy for non-plateau-related neuronal activity) measured in the opto-zone were not significantly affected by ArchT activation (Extended Data Fig. 5c).

Together, these results indicate that OLM interneuron activity constrains BTSP-mediated place cell formation by suppressing BTSP induction in CA1 pyramidal neurons. Silencing OLM neurons during the later phase of learning selectively releases this constraint, allowing additional place fields to form at the reward location, consistent with an increased BTSP occurrence.

### Activating OLM interneurons early in learning suppresses BTSP-mediated place cell formation

To test whether OLM interneuron activity is sufficient to suppress place cell formation, we optogenetically activated Chrna2α-positive OLM neurons during the early phase of spatial learning (Fig. 4, Extended Data Fig. 3g-l). OLM neurons expressing ReaChR^31^ (Chrna2α:ReaChR mice) were activated within the opto-zone during block 1, a phase characterized by high rates of place cell formation and BTSP^5,12^ and low OLM activity (Figs. 1–2). CA1 pyramidal neurons were imaged following AAV-mediated expression of GCaMP6f in Chrna2Δ:ReaChR mice, with identically treated non-ReaChR littermates serving as controls. Activation of OLM interneurons during block 1 significantly inhibited the formation of CA1 spatial representations (Fig. 4a–e; Extended Data Fig. 4d-f). Heatmaps of CA1 population activity revealed a marked reduction in the emergence of place fields within the opto-zone surrounding the reward in ReaChR-expressing animals compared to controls. This suppression was evident both in the overall number of place cells and in their spatial distribution (Fig. 4a-e). Consistent with the block-specific manipulation, place cell development during block 2 was comparable between groups once the optogenetic manipulation was discontinued.

**Fig. 4.**
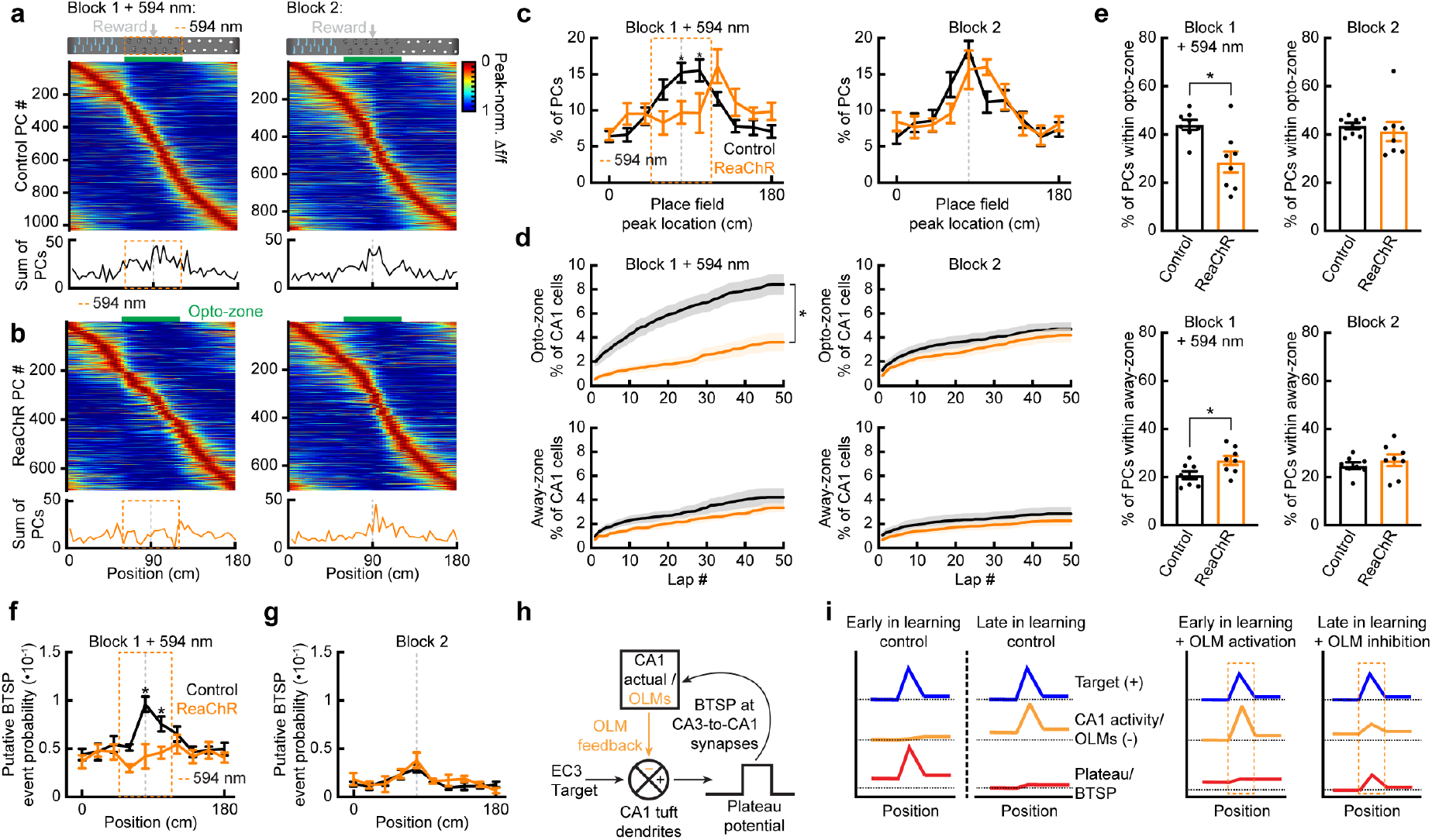
Activating Chrna2Δ-OLMs early in learning. **a**, Top: manipulation paradigm wherein ReaChR in Chrna2Δ-OLMs is activated during block 1 (the opto-zone is 60 cm centered at the reward, the away-zone is the same, but 90 cm away from the opto-zone, orange dashed box depicts 594 nm-light on period). Middle: Blocks 1 and 2 peak-normalized mean Δf/f vs. position for all place cells (PCs) from control mice. Bottom: histogram showing place cell distribution (block 1: n=1031 cells, block 2: n=901 cells). Place cells were analyzed independently within blocks, sorted by place field peak location in the heatmaps. **b**, same as **a** for ReaChR mice (block 1: n=688 cells, block 2: n=702 cells). **c**, Blocks 1 and 2 fractions of CA1 place cells vs. place field peak location (bin=18 cm, two-way RM ANOVA with post-hoc FDR correction, block 1/group x bin: P=1.0•10^-3^, block 1/bin 5: P=7.1•10^-3^, block 1/bin 6: P=4.0•10^-3^). **d**, Blocks 1 and 2 CA1 place cell onsets. Top: place cells with place field peaks within the opto-zone (unpaired two-tailed *t*-test, block 1: P=1.0•10^-3^). Bottom: same as top for away zone. **e**, Place cell distributions in the opto-zone and away-zone. Left: block 1, (unpaired two-tailed *t*-test, opto-zone: P=6.1•10^-3^, away-zone: P=2.8•10^-2^). Right: block 2. **f**, Block 1 mean putative BTSP event probability for all cells vs. position (two-way RM ANOVA, group x bin: P=4.9•10^-3^, bin 5: P<1•10^-4^, bin 6: P=2.3•10^-2^). **g**, same as **f**, but block 2. **h**, BTSP working model with OLM feedback added. **i**, Spatial profile of the proposed signals in early and late learning that regulate BTSP and CA1 spatial representation. Left: typical learning in control animals. Right: manipulation of typical learning by either activation or inhibition of OLMs. In **a**-**g**, control: n=8 mice, ReaChR=8 mice. In **e**, the dots indicate mice. Data are shown as mean ± s.e.m.

We next asked whether this suppression was associated with a reduced probability of observing BTSP. We used the same Ca^2+^ imaging-based criteria as before (Fig. 3h), and found that, in ReaChR-expressing animals, putative BTSP events were significantly reduced during block 1 in the opto-zone (Fig. 4f–g, Extended Data Fig. 5d–e), consistent with OLM activation suppressing BTSP initiation in CA1 pyramidal neurons. We further found that, while control animals showed an approximate 2-fold increase in putative BTSP event probability at the reward, this enrichment was lost in ReaChR animals (1.75±0.12 fold vs. 1.04±0.20 fold for controls vs. ReaChR animals, mean of bins 5–6/mean of bins 1–3, unpaired two-tailed *t*-test P=8.7•10^-3^). Following the end of OLM activation, putative BTSP event occurrence returned to control levels, paralleling the normalization of place field formation. Importantly, this effect was not accompanied by a global change in neuronal activity, as the amplitudes of the lower 80% of Ca^2+^ events were not significantly altered by ReaChR activation (Extended Data Fig. 5f). Together, these experiments demonstrate that increasing OLM interneuron activity early in learning is sufficient to suppress BTSP-mediated place field formation.

## Discussion

Here, we address a central question left open by prior work establishing BTSP as a mechanism for experience-dependent hippocampal learning^9-15^: why BTSP-mediated place field formation declines with learning despite a stable EC3 target signal. We identify a learning-dependent, environment-specific circuit-level inhibitory feedback signal that dynamically regulates when and where BTSP can occur (Fig. 4h–i). This feedback is provided by OLM interneurons, consistent with their circuit position as recipients of CA1 pyramidal cell excitation and inhibitors of distal apical dendrites. Early in learning, low OLM activity permits EC3-driven excitation to trigger dendritic plateaus and robust place field formation, whereas progressive recruitment of OLM-mediated inhibition with learning suppresses further plateau initiation, terminating the development of hippocampal representations once they have converged on their target pattern. Optogenetic manipulations support this model: activating OLM interneurons early suppresses BTSP and place field formation, whereas silencing them later reinstates BTSP and promotes the emergence of additional place fields.

Our findings further support the idea that dendritic plateaus function as locally computed error signals. Rather than relying on a globally broadcast error, plateau initiation in individual CA1 neurons depends on the balance of excitation and OLM-mediated inhibition within the tuft compartment, providing a cell-specific signal that further synaptic modification is warranted^32-34^. An additional feature of this mechanism is its generality across behavioral contexts. We found that OLM activity was biased toward behaviorally salient locations across environments with distinct EC3 target patterns, including one in which salience was defined by a reward-predictive visual cue rather than by reward delivery^12^. Thus, OLM-mediated feedback appears to flexibly reflect task-relevant structure, a property that may be essential for spatial learning in natural environments, where salient features differ across contexts and evolve over time.

Although our results identify OLM interneurons (and specifically the Chrna2*α*-positive subtype) as a core control element of BTSP, additional interneuron classes and neuromodulatory systems are also likely to contribute. Somatic- and dendrite-targeting interneurons^*22,25-28,35-39*^, disinhibitory VIP circuits^40^ activated by novelty^41^, and neuromodulatory inputs^42-44^ have all been implicated in regulating hippocampal plasticity and may interact with OLM-mediated feedback to shape learning^16,40,45^. In hippocampal slices, dendrite-targeting interneurons control dendritic plateau initiation^46,47^, and recent work has shown that a distinct OLM subtype (i.e., NDNF/Nkx2-1-expressing) regulates plateau termination *in vitro*^48^. Determining how these pathways converge to regulate BTSP during behavior remains an important goal for future work.

Together, our findings demonstrate that learning in hippocampal circuits is governed by a dendritic feedback computation. This feedback provides a biologically grounded mechanism for controlling synaptic plasticity at behavioral timescales and across hierarchically organized circuits, which may generalize to other brain regions that require balancing flexibility with stability.

## Acknowledgements

We thank Randy Chitwood and Francisco Mello for technical assistance. We thank Andreas Tolias, Klas Kullander, Simon Chamberland, Richard Tsien, Jun Wu, and Gina Poe for generously providing the SST-IRES-Cre and Chrna2α-Cre mouse lines. We also thank Randy Chitwood, Aaron Milstein, and Sachin Vaidya for valuable discussions. This work was supported by the NIH 4RF1 MH135576 (CG), 1DP2 MH136393 (CG), the Howard Hughes Medical Institute (JCM), and the Cullen Foundation (JCM).

## Author Contributions

EPC, LM, JCM, and CG designed the research. EPC, LM, and CG performed *in vivo* recordings. EPC, LM, JCM, and CG analyzed the data and wrote the manuscript.

## Online methods

All experiments were performed according to methods approved by the Institutional Animal Care and Use Committees at Baylor College of Medicine (Protocol AN-7734) and Brandeis University (Protocols 22022 & 25001).

### Surgery

Experiments were conducted on either sex of adult (older than two months postnatal) SST-IRES-Cre^24^ (n=23, JAX, #018973, also kindly provided by Andreas Tolias), Chrna2Δ-Cre^28^ (n=21, kindly provided by Simon Chamberland, Richard Tsien, Jun Wu, and Gina Poe), and GP5.17^49^ (n=4, JAX, #025393) mice, as well as double transgenic Chrna2α-Cre:GP5.17 (n=14), and Chrna2α-Cre:ReaChR^31^ (JAX, #024846) mice (n=16). Mice were housed in a reverse light cycle facility (12 hours dark/12 hours light) maintained at 30-50% humidity and 21 °C. Surgeries were performed using stereotactic methods while mice were kept under deep anesthesia using isoflurane. Following local antisepsis and the application of a topical anesthetic, the scalp was removed and the skull cleaned. Then, the skull was leveled, and two craniotomy locations for virus injections in dorsal CA1 were marked (2.15 mm | 2.35 mm posterior from bregma, 1.8 mm | 2.2 mm lateral from the midline) as well as the center location of a cranial window (2.2 mm posterior from bregma and 2.0 mm lateral from the midline). Following the virus injection, a 3 mm-diameter craniotomy was made above the hippocampus, and the cortical tissue above the hippocampus was aspirated while rinsed with 0.9% sterile saline. Once the hippocampal capsule was visible, the window implant (a 3 mm-diameter, 1.7 mm-high metal cannula (McMaster) with a window on the bottom and 0.5 mm of shrink tubing around the top) was implanted and cemented in place. Lastly, a titanium head bar was secured to the skull using dental acrylic (Ortho-Jet, Lang Dental). Virus injections were carried out using a microinjector (Drummond, Nanoject II) loaded with a pulled glass pipette that was trimmed and beveled to 25-35 µm and backfilled with mineral oil (Sigma). The virus was loaded by retracting the injector’s plunger. The microinjector was positioned over the craniotomies using a micro-manipulator (MP-285A, Sutter Instruments). For each injection site, a 0.5 mm round craniotomy was drilled, and 46 nl of virus dilution was injected at two depths per site (1200-1300 µm and 1000-1050 µm).

Virus injections varied across experimental paradigms. For imaging SST-OLM axons: AAV2/1.Syn.Flex.GCaMP6f.WPRE.SV40 (Addgene, #100833, titer: 8×10^11^-2×10^12^) with AAV2/1-syn-Flex-TdTomato (UNC Neurotools, titer: 3-9×10^10^). For imaging Chrna2α-OLM somata: AAV2/1.Syn.Flex.GCaMP6f.WPRE.SV40 (titer:8×10^11^-2×10^12^). For inhibition of Chrna2α-OLMs: AAV2/1.hsyn.Flex.ArchT.tdTomato (UNC Neurotools, titer: 3.3×10^11^), or AAV2/1.hsyn.Flex.ArchT.tdtomato (3.3×10^11^) with AAV2/1.Syn.GCaMP6f.WPRE.SV40 (Addgene, #100837, titer: 1-3×10^12^). For inhibition controls: AAV2/1-syn-Flex-tdTomato (titer: 3×10^9^-9×10^10^), or AAV2/1-syn-Flex-tdTomato (titer: 3×10^9^-9×10^10^) with AAV2/1.Syn.GCaMP6f.WPRE.SV40 (titer: 1-3×10^12^). For activation of Chrna2α-OLMs and controls: AAV2/1.Syn.GCaMP6f.WPRE.SV40 (Janelia Viral Core, titer:1×10^12^).

### Behavioral training and task on the linear track treadmill

The treadmill comprised a fabric belt (180 cm; McMaster Carr) and a custom-built lick port that dispensed sucrose water, controlled by a solenoid valve (LHQA1231220H, The Lee Company). The belt was self-propelled by the animal, and its speed was tracked using an encoder (Broadcom) at the front wheel axle. An Arduino-based behavioral state machine (Bpod, Sanworks) and corresponding MATLAB GUI were used to control the reward value and sensors, and to track the encoder. Light delivery for optogenetic manipulations was controlled by a Teensy 3.5 microcontroller interfaced with a custom-written MATLAB GUI, or by the behavioral state machine and pulse train generator (PulsePal, Sanworks). The behavioral data, stimulations, and frame times were monitored and recorded using a NIDAQ-based acquisition system (National Instruments, PCIe-6341) and the MATLAB-based Wavesurfer software (version 0.982, Janelia).

5 to 7 days following surgery, the mice were placed on a water restriction protocol (receiving 1.5 ml per day). Three or four days into water restriction, animals were habituated to the experimenter for 5-6 days, followed by ∼3-5 days of head-fixed treadmill training. Training days were conducted during the animal’s dark cycle on a belt with no cues and with a sucrose reward (5-10% solution) dispensed at a random location each lap. For mice later used in optogenetic experiments, 594 nm light was delivered at random locations through the objective during the training to habituate animals to the presence of light. During experimental days for the tactile cue paradigm, mice learned to navigate a cue-rich belt (three cue sections: Velcro tape patches, thin hot glue sticks, and correction fluid dots) with a single fixed location for a sucrose reward. In the light-cue learning experiments (Extended Data Fig. 2), the animals were on a blank belt (no tactile cues) and received a bilateral visual stimulus of blue light (10 Hz for 500 ms) 50 cm before the fixed reward location. For OLM activity recording experiments, recordings lasted 30 to 60 minutes. Mice were excluded if they ran <75 laps or >350 laps in 60 minutes. Mice recorded past day 1 (Extended Data Fig. 1) were recorded for a maximum of three more days (up to day 4). For the optogenetic experiments, analyses were limited to 88-120 laps (mean±SEM: 101±1 laps) per session. Animals were recorded at most once per day.

### In vivo two-photon Ca^2+^ imaging

All experiments were conducted in a dark box, and Ca^2+^ images were captured using custom-built two-photon microscopes (detailed descriptions below). Experiments recording OLM interneuron activity were performed on both microscopes. Optogenetic experiments activating ArchT or ReachR were performed on the microscopes at Brandeis or Baylor, respectively. The images were recorded using ScanImage at 30 Hz and 512 x 512 pixels. Fields of view were 280 x 280 um and for animals with multiple recording days, we attempted to return to the same field of view (Extended Data Fig. 1a). GCaMP6f, and in some cases tdTomato or Citrine, were imaged through Nikon 16x, 0.8-numerical-aperture objectives (both microscopes) and excited at 920 nm (typically 30-75 mW measured under the objective)

For experiments conducted at Baylor recording the activity of CA1 OLM interneurons, the microscope (Janelia MIMMS 2.0 design) used a Ti:Sapphire laser (Coherent, Chameleon Ultra II), passed through a FF705-Di01 (Semrock) primary dichroic mirror. Images were collected with ScanImage R2022 premium software (Vidrio). Emission light passed through a 565 DCXR dichroic mirror (Chroma) and either a 531/46 nm (GCaMP6f channel; Semrock) or a 612/69 nm (tdTomato channel, Semrock) bandpass filter. Light was detected by two GaAsP photomultiplier tubes (11706P-40SEL, Hamamatsu). For optogenetic ReaChR manipulation experiments, in which only the GCaMP6f channel was recorded, a 594 nm laser (Obis, Coherent) was used. Diffuse 594 nm light was reflected into the beam path by a DMLP805R (Semrock) dichroic mirror, the filter in front of the PMT in the GCaMP6f channel was replaced with stacked 505/19 nm (Semrock) and 514/44 nm (Semrock) bandpass filters, and the primary dichroic was substituted with a Di02-R561 (Semrock) dichroic mirror.

For experiments conducted at Brandeis, the microscope (INSS, UK) included a Spectra-Physics Insight X3 laser. Images were collected with ScanImage R2021 software (Vidrio). Emission light passed through a T565lpxr dichroic mirror (Chroma) and either an ET 510/80 nm (GCaMP6f channel, Chroma) or a 630/75 nm (tdTomato channel, Chroma) bandpass filter, then was detected by two GaAsP photomultiplier tubes (11706P-40, Hamamatsu). For optogenetic ArchT manipulation experiments, in which only the GCaMP6f channel was recorded, diffuse 594 nm light from an LED (Thorlabs) passed through an ET570LP low-pass filter (Chroma), and the filter in front of the PMT in the GCaMP6f channel was replaced with an ET 510/80 (Chroma) bandpass filter stacked between two BG39 (Chroma) filters.

### Optogenetic perturbation of OLM interneurons in CA1

To examine the effect of OLM inhibition (experiments performed at Brandeis), we used the following groups: 1) experimental mice (Chrna2α-Cre:GP5.17 injected with AAV2/1.hsyn.Flex.ArchT.tdtomato, n=6; Chrna2α-Cre injected with AAV2/1.hsyn.Flex.ArchT.tdtomato and AAV2/1.Syn.GCaMP6f.WPRE.SV40, n=3); 2) control mice (Chrna2α-Cre:GP5.17 injected with AAV2/1.hsyn.Flex.tdtomato, n=8; Chrna2α-Cre injected with AAV2/1.hsyn.Flex.tdtomato and AAV2/1.Syn.GCaMP6f.WPRE.SV40, n=2; non-injected GP5.17, n=4). The experimental (ArchT-expressing) and control mice underwent the same procedures and paradigm: the window was implanted in the same surgery as the injections. Imaging and optogenetic experiments were conducted 18 to 28 days post-surgery. Recordings were obtained on days 1–3 on the cue-enriched, fixed-reward belt. Continuous diffuse light of 594 nm from an LED (Thorlabs) at 4 mW measured under the objective was delivered through the objective. In all optogenetic experiments, the region of the belt where 594 nm light was applied (the “opto-zone”) was 60 cm wide, centered at the reward location. Light was applied only in block 2 (approximately 50 laps). To validate the effect of ArchT activation on OLM activity (Extended Data Fig. 3a-f), we recorded OLM activity in a subset of Chrna2α-Cre mice that expressed GCaMP6f and ArchT (n=3) or GCaMP6f and tdT (n=2) on the final day of experimentation. A field of view overlying the pyramidal neuron imaging plane was selected that contained OLM neurons expressing GCaMP6f together with either ArchT or tdTomato (Chrna2α-positive cells), as well as neurons expressing only GCaMP6f (Chrna2α-negative cells). Ca^2+^ signals were recorded while the LED was on for approximately 2.5-3 seconds, or, in some laps, for 60 cm of the animal’s travel on the belt.

To examine the effect of OLM excitation (experiments performed at Baylor), we used the following groups: 1) experimental mice (Chrna2α-Cre:ReaChR injected with AAV2/1.Syn.GCaMP6f.WPRE.SV40, n=8); 2) littermate control mice (Chrna2a-Cre:ReaChR litter mates that were confirmed by genotype to lack at least one allele required for ReaChR expression, injected with AAV2/1.SynGCaMP6f.WPRE.SV40, n=8). The experimental ReaChR-expressing mice and control mice underwent the same procedures and paradigm as ArchT mice in OLM inhibition experiments. Optogenetic manipulation in ReaChR mice was restricted to day 1 on the cue-enriched, fixed-reward belt. Diffuse light of 594 nm was delivered from a laser (Obis, Coherent) at 40 Hz (square pattern) at 0.8-0.95 mW measured under the objective. 594 nm light was only applied during block 1 (50 laps) while the mouse was within the opto-zone. The opto light was not applied if the mouse’s velocity dropped below 2.5 cm/s, or if it had been applied for more than 5s continuously. To validate the effect of ReaChR activation on OLM activity (Extended Data Fig. 3g-l), stratum oriens interneurons were found directly above the field of view used for pyramidal neuron recordings on the last day of experimentation. Ca^2+^ signals were recorded while the laser was either activated for 5 seconds (“fixed-duration”, n=8 mice) or within the opto-zone (“fixed-location”, n=7 mice). Each mouse was validated to have responsive OLMs, which showed > 2-fold increases in mean Δf/f while the laser was on (for fixed-duration: a 3s window before stimulation was compared with a 4s window during; for fixed-location: the opto-zone was compared with surrounding locations).

### Data analysis

#### Ca^2+^ signal extraction and activity map generation

To extract axonal Ca^2+^ signals from the population of CA1 SST-expressing OLM interneurons, motion correction was performed using Suite2p^50^ (version 0.14.4), with both green (GCaMP6f) and red (tdTomato) channels aligned, and the red channel used for image registration. GCaMP6f fluorescence was then averaged across the field of view using custom MATLAB functions (version 2024b), excluding a narrow border (2% of frame width) to minimize motion-related artifacts. For OLM somatic Ca^2+^ imaging, motion correction and functional region of interest (ROI) detection were performed using Suite2p. For pyramidal neuron recordings from the ArchT mice, ROIs were identified applying a custom-pretrained Cellpose^51^ (version 2.3.2) model in Suite2p. In ReaChR mice, which exhibited dense pyramidal neuron labeling, functional ROI detection was used and cross-validated with the same Cellpose model. All ROIs were manually curated, and those with insufficient signal quality were excluded. Interneurons were identified based on event amplitude, duration, and frequency, and manually removed. All datasets in which motion correction was unsuccessful were excluded.

Further analyses were performed using custom MATLAB code. First, raw fluorescence signals were converted to Δf/f, calculated as (f − f_0_)/f_0_. For pyramidal neurons, f_0_ was defined as the mode of the fluorescence histogram, whereas for OLM interneurons, f_0_ was defined as the 8th percentile of fluorescence values. For pyramidal cell recordings, this was followed by neuropil correction (subtraction of 70% of the neuropil Δf/f signal obtained via Suite2p). Significant Ca^2+^ events were identified as transients exceeding three standard deviations of the baseline noise (mode of the histogram).

For all recordings, spatial maps were next generated by dividing the 180-cm belt (one lap) into 50 spatial bins (3.6 cm each) and taking the mean of αf/f values from periods of running (when the animal’s speed exceeded 2.5 cm/s) within each spatial bin for every lap. For pyramidal neuron recordings, where cellular density can lead to cross-contamination, these spatially binned activity maps were used to calculate noise correlations between neighboring ROIs (within three soma diameters). In this noise correlation, Pearson’s correlation coefficient thresholds of 0.35–0.5 were applied to identify potential signal cross-contamination, and contaminated ROIs were excluded from further analysis. For recordings of OLM activity in learning, spatial αf/f maps were multiplied by the animal’s dwell time within each spatial bin. Dwell time correction was used to account for learning-related changes in running speed and isolate spatial patterns of inhibitory activity, as OLM interneurons have been demonstrated to be highly velocity modulated^25-27^. Dwell time was computed as d^bin^/V^bin^, where d^bin^ is the spatial bin width (3.6 cm), and V^bin^ is the mean velocity within that bin. Dwell-time correction reduces speed-related distortion but does not remove effects of licking or reward timing. To compute the population-averaged somatic activity for recordings of Chrna2α-expressing OLMs, the mean Δf/f•s was calculated across the spatially binned activity maps of each cell recorded per mouse. To quantify pooled pairwise correlations from Chrna2α-expressing somata recorded together (Fig. 2o), linear activity vectors were constructed from spatial bins within lap quintiles for each neuron and Pearson’s correlation coefficients were computed for each unique pair of neurons per mouse. For spatial display of the mean z-scored Δf/f•s activity across groups of mice, spatial maps from each mouse were linearly duplicated by interpolation along the laps dimension to 1000 laps (each lap representing 0.1% of total laps run; these duplicated spatial maps were used for presentation only and not quantified in any way). For visualization, spatial maps were smoothed using a three-point boxcar filter and centered such that either the reward location (cue-belt environment) or the visual cue location (visual cue environment) corresponded to spatial bin 25.

#### CA1 place cell identification

Place cells were determined as previously described^12,20^. In brief, from each CA1 pyramidal neuron’s spatial map of mean Δf/f, we first identified potential onset laps for a place field(induction lap). Place fields were defined as those spatial bins with contiguous activity >20% of the peak mean Δf/f. Induction occurred at a lap where significant Ca^2+^ activity was within the neuron’s eventual place field, and two out of the following five laps also had significant activity in the eventual place field. If multiple laps fit these criteria per neuron, the first instance was the induction unless the place field generated was weak and disappeared for more than 20 laps at some point. Only the laps following place field induction were used to determine place cell identity. CA1 place cells were defined as neurons exhibiting (1) significant spatial information about the linear track position (outside the 95% confidence interval of values obtained from shuffled data) and (2) high place field reliability, defined as significant activity on more than 33% of laps following place field induction. The spatial information was calculated as described previously^20^ and compared to 100 shuffles of the activity. For shuffling, Δf/f from periods when the mouse’s velocity exceeded 2.5 cm/s was circularly rotated 500 frames and divided evenly into six sections that were permuted in pseudo-random order. Place cells were determined independently across the laps of blocks 1 and 2, within which only one place field was used per neuron. If the spatial bin containing the peak of a place cell’s place field moved by >20 cm between the laps of blocks 1 and 2, its block 2 place field was considered a new place field for that cell.

#### Putative BTSP event detection

Putative BTSP event detection (Figures 3-4) was performed across all cells and all spatial locations. Only cells that had events with amplitude >1Δf/f and at least 20 significant Ca^2+^ events (amplitude greater than three times the standard deviation of baseline noise) were included in this analysis (these were considered “active neurons”). Putative BTSP events were identified using criteria adapted from whole-cell recordings and previous descriptions using GCaMP6f imaging^5,9,12,30^. These were events that (1) were large Ca^2+^ transients, reaching amplitudes within the top 10% of all significant events for each neuron, and >1 Δf/f; (2) were followed by significant activity within 45 cm of the event peak in at least three of the subsequent five laps; (3) were followed by a forward shift (relative to the event peak) in the mean Δf/f center of mass across the subsequent five laps; (4) Only the first event meeting the criteria (1)-(3) was included in subsequent analyses. Spatial maps of putative BTSP events were generated for each cell, wherein every spatial bin x lap was scored as having a putative BTSP onset (1) or not (0). The putative BTSP event onset was the time at which the event exceeded the significance threshold of 3 standard deviations above the noise band. Putative BTSP event probability per animal for blocks 1 and 2 was calculated as the total number of detected putative BTSP event onsets within each spatial bin (10 bins, 18 cm per bin), normalized by the number of ROIs analyzed in the animal. For determining the relationship between the induction velocity and the place field width, the mean velocity (cm/s) within the spatial bin that contained the peak of the putative BTSP event was calculated and plotted as a function of the place field width computed across the five to ten laps following the putative BTSP event.

#### Behavioral data quantification

As with the analysis of Ca^2+^ data, spatial maps for behavioral data were constructed by first dividing the belt length (also one lap, 180 cm) into 50 spatial bins (3.6 cm each). Along all laps, the mean velocity was averaged within each spatial bin. Lick probability was computed by scoring each spatial bin as containing (1) or lacking (0) at least one lick by the mouse, then by taking the mean along this binary matrix.

#### Statistical methods

The exact sample size (n) for each experimental group is indicated in the figure legend or in the main text. No statistical methods were used to predetermine sample sizes, but our sample sizes are similar to those reported in previous publications. In some cases, when the data distribution was assumed but not formally tested to be normal, data were analyzed using two-tailed paired or unpaired *t*-tests or one-way or two-way RM ANOVAs with post-hoc FDR correction, as stated in the text or figure legends. Data were analyzed automatically without consideration of trial conditions or experimental groups. Experiments and data analyses were not randomized and not performed blind to the experimental conditions. If not otherwise indicated in the figure, data are shown as mean ± SEM.

## Data Availability

The data that support the findings of this study are available from the corresponding author upon request.

## Code Availability

The code that supports the findings of this study is available from the corresponding author upon request.

