## Supplemental Information for "Learning-dependent feedback by OLM interneurons shapes CA1 representations"

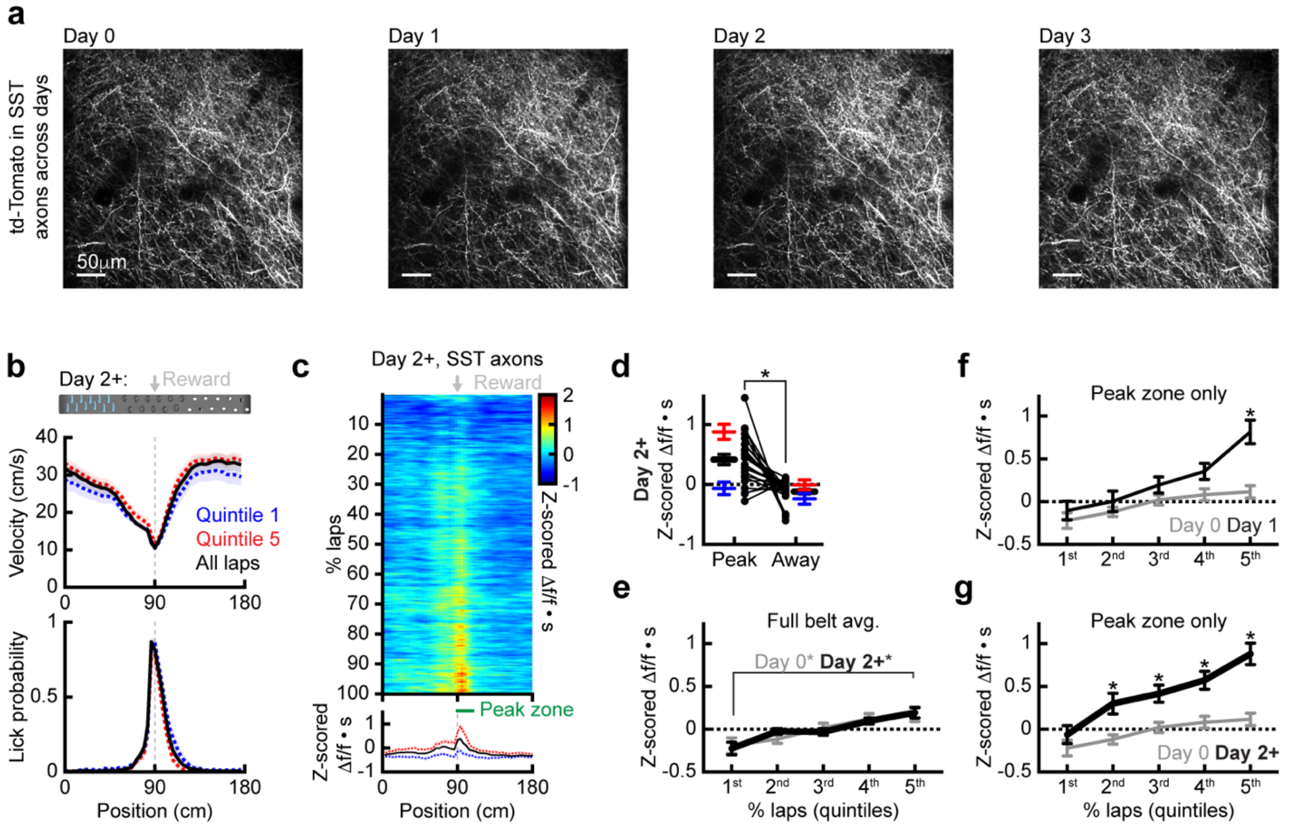


**Extended Data Fig. 1 OLM activity during subsequent days of learning. a**, Cross-day representative, two-photon, time-averaged images of SST-OLM axons expressing tdTomato in CA1 SLM. **b**, Pooled recordings from day 2 and subsequent days (day 2+, see methods) of SST-OLM axons for mice on a belt with tactile cues and one fixed reward (n=23 recording sessions). Top: task design (same as **Fig. 1**). Middle: mean running profile across space. Bottom: mean licking probability across space. **c**, Days 2+ mean z-scored Δf/f•s. Top: along the percent of laps run. Bottom: spatial profiles (blue: quintile 1, red: quintile 5, black: all laps). The peak-zone is 18 cm beginning at the spatial bin with reward, the away-zone is the same but 90 cm away. **d**, Mean z-scored Δf/f•s within the peak-zone and away-zone for day 2+ (blue: quintile 1, red: quintile 5, black: all laps) (paired two-tailed *t*-test, $P=2 \cdot 10^{-4}$). **e**, Mean z-scored Δf/f•s vs. lap quintiles for day 0 and day 2+ recordings (two-way RM ANOVA with post-hoc FDR correction, day 0/quintile 1 vs. 5: $P=2.1 \cdot 10^{-3}$, day 2+/quintile 1 vs. 5: $P<1 \cdot 10^{-4}$). **f**, Mean z-scored Δf/f•s within peak zone only vs. lap quintiles for day 0 (gray) vs. day 1 (thin, black) (two-way RM ANOVA with post-hoc FDR correction, group x quintile: $P=5.5 \cdot 10^{-3}$, quintile 5: $P<1 \cdot 10^{-4}$). **g**, Mean z-scored Δf/f•s within peak zone only vs. lap quintiles for day 0 (gray) vs. day 2+ (thick, black) (two-way RM ANOVA with post-hoc FDR correction, group x quintile: $P=8.1 \cdot 10^{-3}$, quintile 2: $P=5.8 \cdot 10^{-3}$, quintile 3: $P=9.3 \cdot 10^{-3}$, quintile 4: $P=1.3 \cdot 10^{-3}$, quintile 5: $P<1 \cdot 10^{-4}$). In **d**, the dots represent mice. Data are shown as mean ± s.e.m.

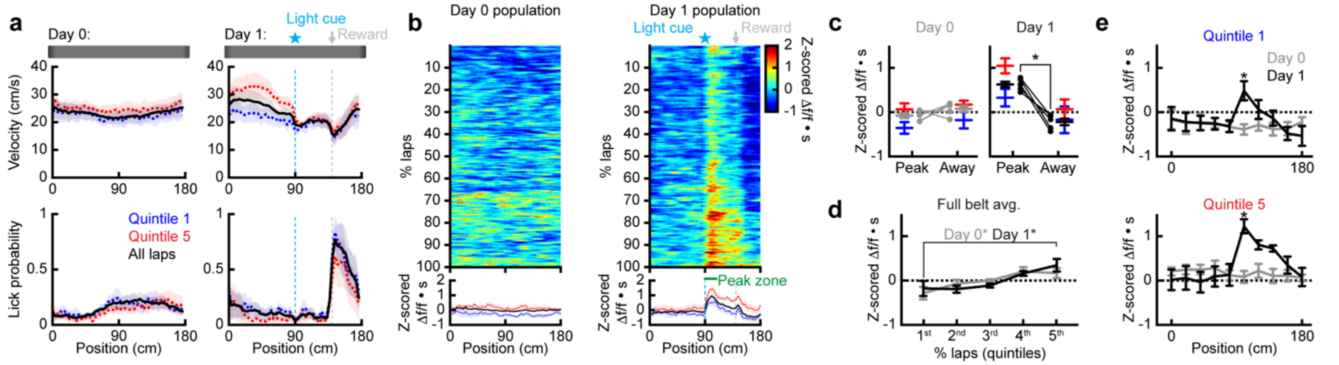


**Extended Data Fig. 2 OLM activity from mice in a light cue learning task. a**, Top: task design of day 0 (the last day of habituation), and day 1 (the first day of learning on a belt with a light cue). Middle: running profile across space. Bottom: licking probability across space. **b**, Days 0 and 1 mean z-scored Δf/f•s. Top: along the percentage of laps run. Bottom: spatial profiles (blue: quintile 1, red: quintile 5, black: all laps). The peak-zone is 18 cm beginning at the spatial bin with the light cue, and the away-zone is the same, but 90 cm away. **c**, Mean z-scored Δf/f•s within the peak-zone and away-zone (blue: quintile 1, red: quintile 5, black/gray: all laps). Left: day 0. Right: day 1 (paired two-tailed *t*-test, $P=1.5 \cdot 10^{-3}$). **d**, Mean z-scored Δf/f•s vs. lap quintile (two-way RM ANOVA with post-hoc FDR correction, day 0/quintile 1 vs. 5: $P=1.1 \cdot 10^{-2}$, day 1/quintile 1 vs. 5: $P=3.6 \cdot 10^{-3}$). **e**, Quintiles 1 and 5 mean z-scored Δf/f•s vs. position (bin=18 cm**,** two-way RM ANOVA with post-hoc FDR correction, quintile 1/group x bin: $P<1 \cdot 10^{-4}$, quintile 1/bin 6: $P=7 \cdot 10^{-4}$, quintile5 group x bin: $P<1 \cdot 10^{-4}$, quintile 5/bin 6: $P<1 \cdot 10^{-4}$). For days 0 and 1, n=5 mice. In **c**, the dots represent mice. Data are shown as mean ± s.e.m.

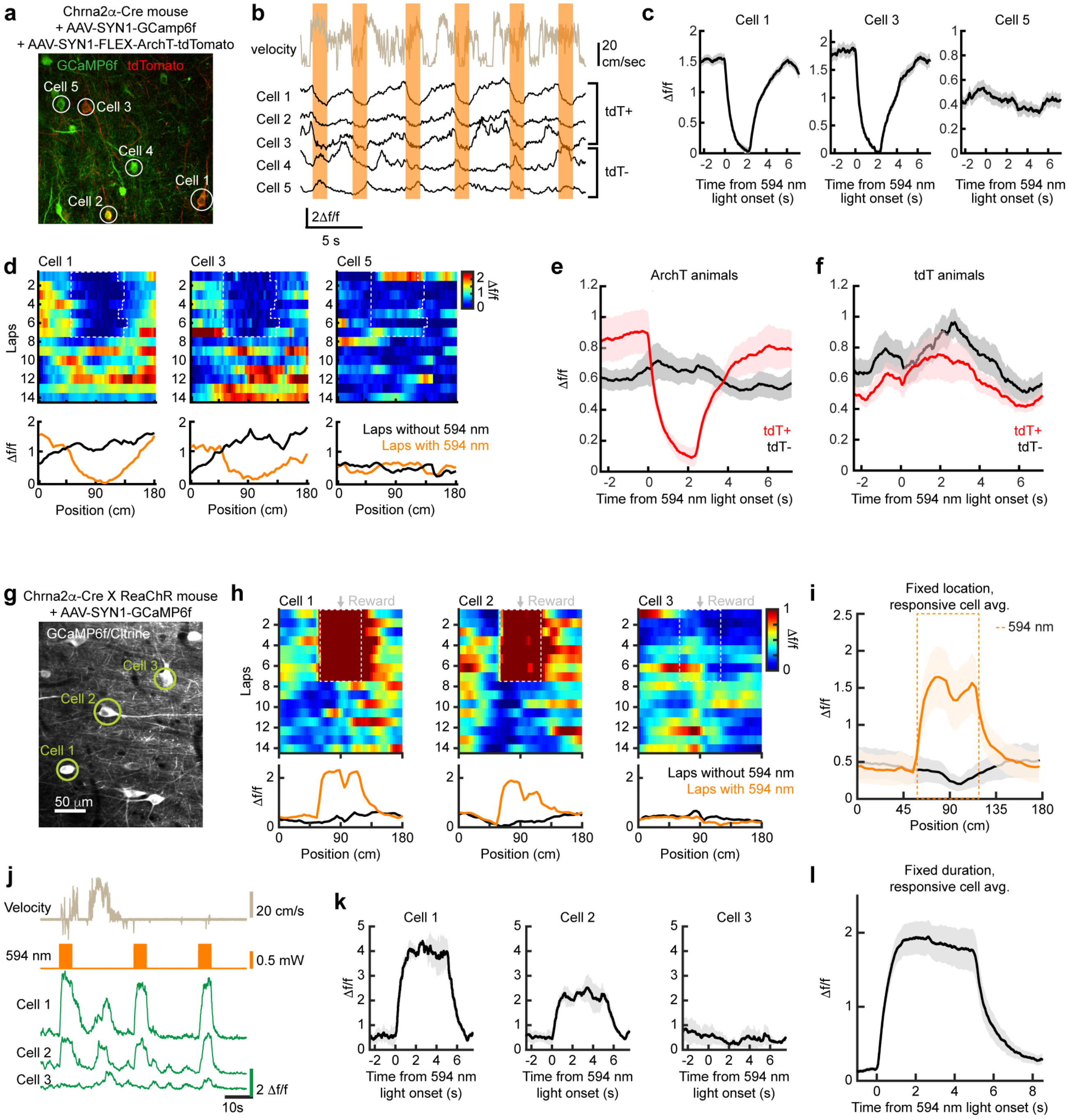


**Extended Data Fig. 3 Optogenetics validation of Chrna2α-OLMs.** **a**, Representative, two-photon, time-averaged image of stratum oriens interneurons with ArchT-tdTomato (red) expressed in Chrna2α-OLMs and pan-neuronal expression of GCaMP6f (green). Neurons indicated correspond to **b**-**d**. **b**, Example traces of velocity, and GCaMP6f Δf/f for three ArchT-expressing OLMs and two non-ArchT-expressing neurons. Orange bars indicate 594 nm-light on. **c**, Mean Δf/f vs. time relative to the onset of an approximately 2.5 second-long 594 nm-light pulse (n=6 trials). **d**, Mean Δf/f. Top: along laps run. The white dashed frame indicates 594 nm-light on. Bottom: spatial profiles. **e**, Mean Δf/f vs. time relative to 594 nm-light onset in ArchT-tdTomato-expressing mice (n=3 mice, ArchT-tdTomato positive neurons: n=11; ArchT-tdTomato negative neurons: n=14). **f**, Same as **e**, but tdTomato-expressing mice (n=2 mice, tdTomato positive neurons: n=9. tdTomato-negative neurons: n=11). **g**, Representative, two-photon, time-averaged image of stratum oriens interneurons with ReaChR-Citrine expressed in Chrna2α-OLMs and pan-neuronal expression of GCaMP6f. Neurons indicated correspond to **h**,**j**,**k**. **h**, Mean Δf/f for two responsive neurons and one non-responsive. Top: along laps run. Bottom: spatial profiles. **i**, Mean Δf/f vs. position for laps with or without 594 nm-light (mice: n=7, cells, n=21). **j**, Example traces of velocity, approximate 594 nm-light intensity (5 second pulses of 40 Hz), and GCaMP6f Δf/f for three neurons. **k**, Mean Δf/f vs. time relative to the 594 nm-light onset (n=3 trials). **l**, Mean Δf/f vs. time relative to the 594 nm-light onset (mice: n=8, cells: n=34). Mice used for the data in **g-l** are the same individuals used for the data in **Fig. 4**. Data are shown as mean ± s.e.m.

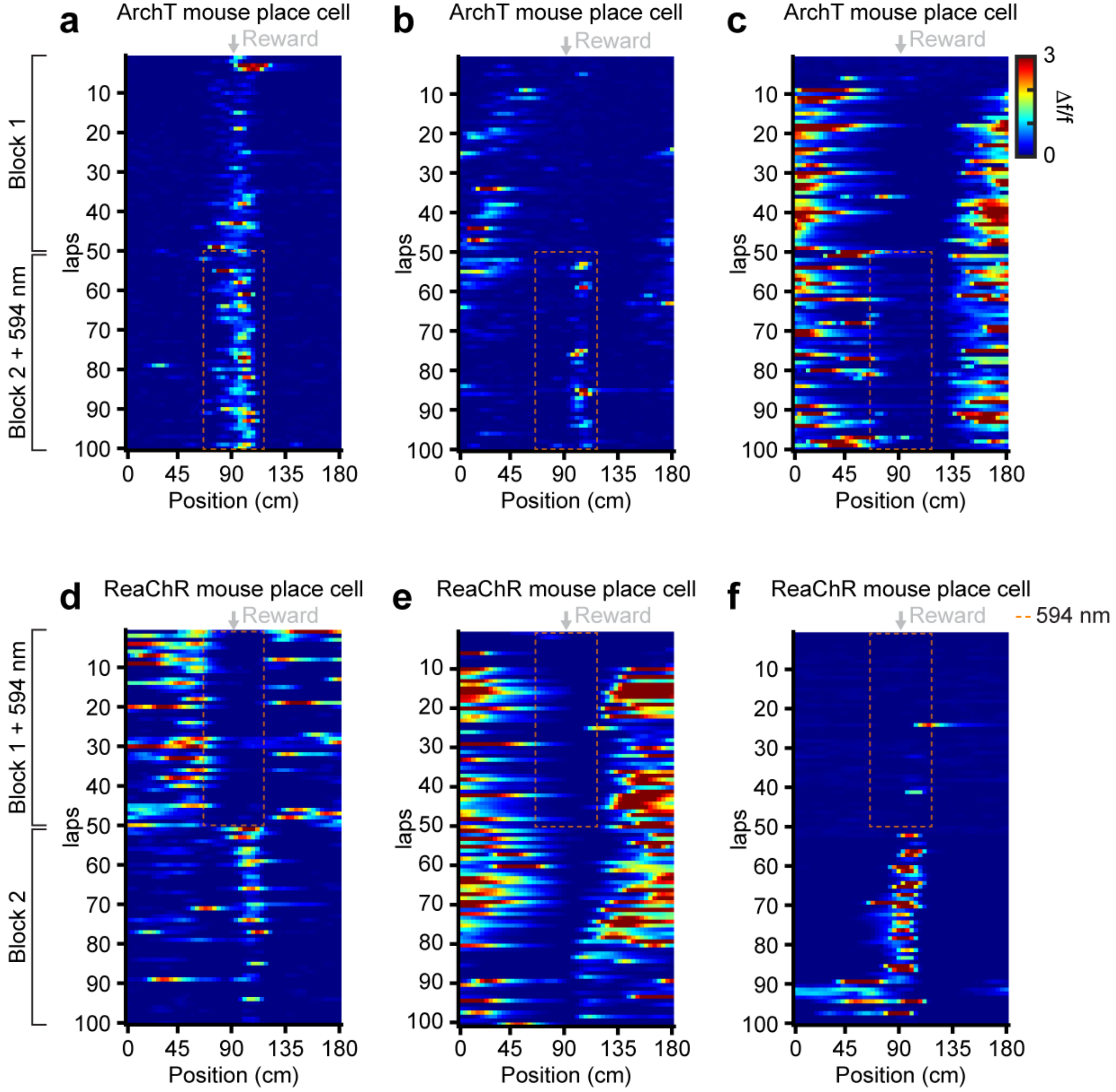


**Extended Data Fig. 4 Example place cells from opsin-expressing Chrna2a mice. a-f**, Example place cells across blocks 1 and 2 (50 lap blocks with 594 nm light provided for optogenetic manipulation as indicated by orange dashed box). Mean Δf/f along laps. **a-c** Place cells from ArchT mice. **d-f** Place cells from ReaChR mice.

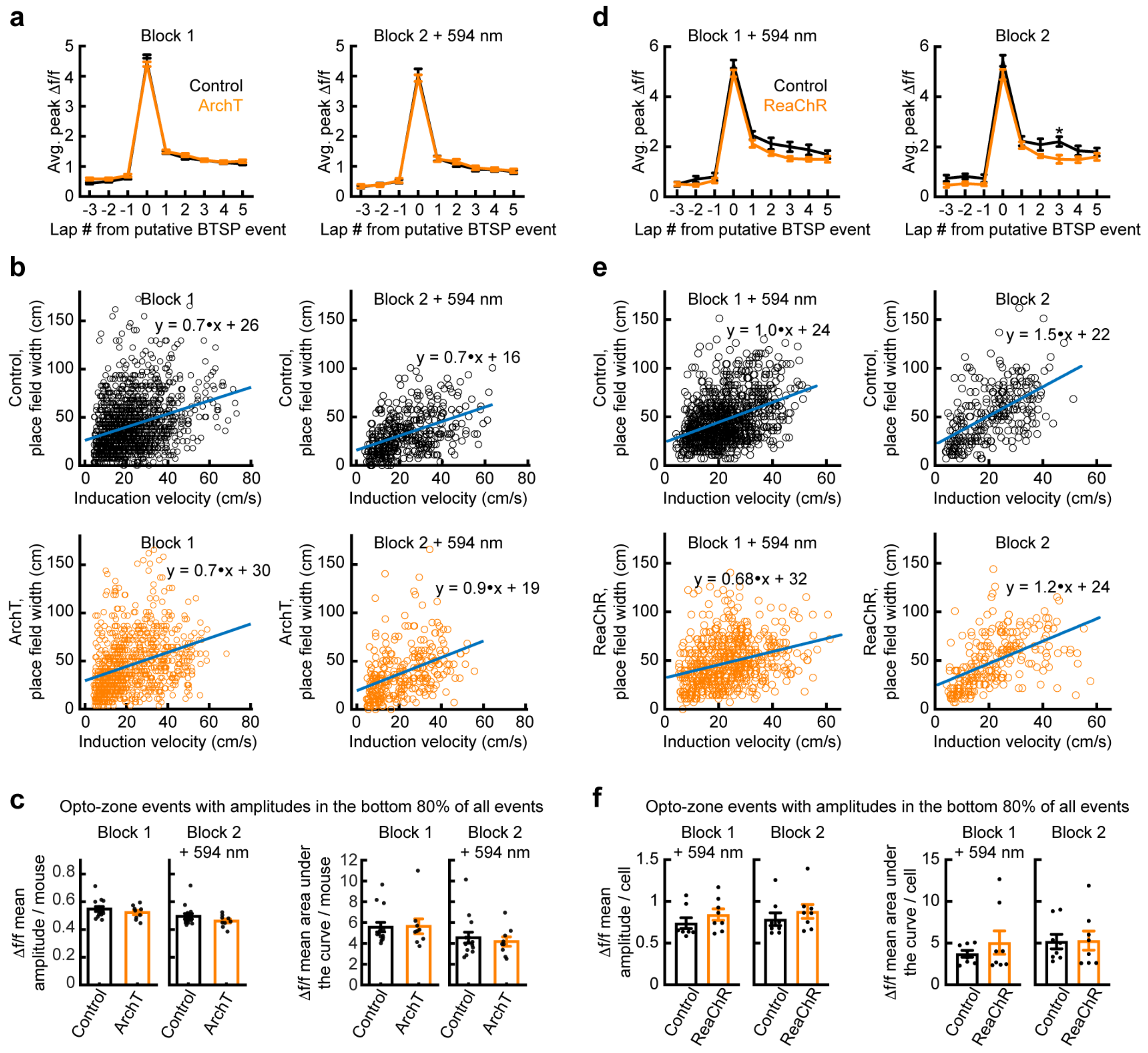


**Extended Data Fig. 5 Further pyramidal neuron events in opsin-expressing Chrna2α mice. a-c**, Events from mice with ArchT expressed in Chrna2α–OLMs. **a**,**b** BTSP signatures in CA1 place cells. Left: for place fields formed in block 1 (control mice: n=1495 events, ArchT mice: 891 events). Right: newly formed place cells in block 2 (control mice: n=398 events, ArchT mice: 300 events). **a,** The abrupt establishment of place fields following putative BTSP events. Mean peak Δf/f vs. lap (two-way RM ANOVA with post-hoc FDR correction, group x lap: not significant). **b**, Place field width (from within five to ten laps following the putative BTSP event) vs. the velocity at the putative BTSP event. Top: Control. Bottom: ArchT. **c**, The effect of Chrna2α-OLM modulation on non-plateau events for all cells from blocks 1 and 2. Left two: mean Δf/f amplitude. Right two: mean Δf/f area under the curve. For panels **a-c**, control: n=14 mice, ArchT-expressing: n=9 mice. Mice used for data in **a-c** are the same individuals used for data if **Fig. 3**. **d-f**, same as panels **a-c** but for mice with ReaChR expressed in Chrna2α–OLMs. **d, e**, Block 1 control mice: n=1025 events, block 1 ReaChR mice: n=636 events, block 2 control mice: n=276 events, block 2 ReaChR mice: n=241 events. **d**, block 2: two-way RM ANOVA with post-hoc FDR correction, group x lap: not significant, lap 3 from putative BTSP event: $P=1.7\cdot10^{-3}$. For panels **d-f**, control: n=8 mice, ReaChR n=8 mice. Mice used for data in **d-f** are the same individuals used for data in **Fig. 4**. Data are shown as mean ± s.e.m.
